# Development of SARS-CoV-2 Nucleocapsid Specific Monoclonal Antibodies

**DOI:** 10.1101/2020.09.03.280370

**Authors:** James S. Terry, Loran BR Anderson, Michael S. Scherman, Carley E. McAlister, Rushika Perera, Tony Schountz, Brian J. Geiss

## Abstract

The global COVID-19 pandemic has caused massive disruptions in every society around the world. To help fight COVID-19, new molecular tools specifically targeting critical components of the causative agent of COVID-19, SARS-Coronavirus-2 (SARS-CoV-2), are desperately needed. The SARS-CoV-2 nucleocapsid protein is a major component of the viral replication processes, integral to viral particle assembly, and is a major diagnostic marker for infection and immune protection. Currently available antibody reagents targeting the nucleocapsid protein were primarily developed against the related SARS-CoV virus and are not specific to SARS-CoV-2 nucleocapsid protein. Therefore, in this work we developed and characterized a series of new mouse monoclonal antibodies against the SARS-CoV-2 nucleocapsid protein. The anti-nucleocapsid monoclonal antibodies were tested in ELISA, western blot, and immunofluorescence analyses. The variable regions from the heavy and light chains from five select clones were cloned and sequenced, and preliminary epitope mapping of the sequenced clones was performed. Overall, the new antibody reagents described here will be of significant value in the fight against COVID-19.

## Introduction

Over the course of the last nine months, the novel SARS-CoV-2 coronavirus has spread dramatically across the world, causing the severe respiratory illness termed COVID-19. There have been over 25 million reported cases of COVID-19 globally as of August 2020 (1), and over 845 thousand reported deaths attributed to this devastating disease. SARS-CoV-2 is a respiratory droplet-borne pathogen (2) and is easily transmitted between individuals in close proximity, leading to explosive spread and a dire need for rapid diagnostic testing to help control outbreaks.

Testing for COVID-19 infection currently focuses primarily on detection of viral genomic RNA present in patient respiratory samples, including nasopharyngeal swabs and nasal samples. Because COVID-19 is a respiratory disease, detection of viral genomic RNA in patient nasal samples is a positive indicator of both infection and the potential for an infected individual to spread the virus to others. The current diagnostic for detecting viral genomic RNA is quantitative reverse-transcriptase polymerase chain reaction (qRT-PCR), which can sensitively detect the presence of viral RNA in samples (3–5) and can be automated for to test large numbers of samples in parallel. This workhorse assay can provide exquisitely sensitive and specific detection of SARS-CoV-2 infection, but faces challenges. Those challenges include significant pre-processing of samples such as RNA extraction, high cost of reverse-transcription quantitative PCR reagents, and the need for sophisticated real-time capable thermocyclers for performing the PCR procedure (6). Additionally, RNA is only one of a number of analytes that can provide significant clinical value for diagnosing infection. The coronavirus nucleocapsid protein is one such analyte.

Coronavirus RNA genomes are coated with nucleocapsid protein within viral particles and within infected cells. The nucleocapsid (N) protein is a ∼50kDa protein that forms dimers that oligomerize on viral RNA, providing protection of the viral genome from cellular RNA decay enzymes and compacting the viral genome into a small enough package to fit within virion particles (7–10). There have been estimates that between 720 and 2200 nucleocapsid monomers are present for every viral RNA genome copy within virion particles (10–15), making the nucleocapsid protein an intriguing analyte for viral infection. Several publications from the original SARS-CoV outbreak in 2003-2004 indicated that detection of nucleocapsid in patient serum samples is diagnostic for early SARS disease, and the amount of detectable SARS-CoV nucleocapsid antigen present in patient samples tracked well with viremia (16–20). More recent data from the SARS-CoV-2 pandemic indicate that N protein is found in very low but detectable amounts in patient serum (21), but N protein has been found in greater amounts in patient nasopharyngeal swab and anterior nares swab samples.(22) Given the high copy number of the N protein compared to viral genomes and the relative stability of N protein in patient samples, detection of N can serve as a valuable orthogonal diagnostic marker compared to genome detection by RT-qPCR. Detection of protein analytes requires specific antibodies, and since SARS-CoV-2 has emerged very recently, no SARS-CoV-2 specific antibodies have been reported in the literature. There is significant homology between SARS-CoV and SARS-CoV-2, new antibodies need to be produced for the research community that may have increased specificity and utility for detecting SARS-CoV-2 nucleocapsid protein or for potential therapeutic use (23–25).

Here we report the generation and characterization of a panel of monoclonal antibodies targeting the SARS-CoV-2 N protein. We expressed and purified a truncated recombinant N protein, used the recombinant antigen to immunize mice and generated a panel of hybridomas, and tested the resulting clones for activity in western blots, ELISAs, and immunofluorescence assays with SARS-CoV-2 infected cells. Cross-reactivity of the antibodies against SARS-CoV, HuCoV-NL63, and HuCoV-229E N protein was tested. We determined the V_H_ and V_L_ sequences of the top 5 clones and performed epitope mapping to identify antigenic regions within the N protein. Overall, our data provides a strong foundation for using these monoclonal antibodies to study SARS-CoV-2 N protein and development of novel diagnostic assays to detection of COVID-19.

## Materials and Methods

### Expression and purification of Coronavirus N proteins

Amino acid sequences for coronavirus N proteins (SARS-CoV-2 (YP_009724397.2), SARS-CoV (ABI96968.1), MERS (YP_009047211.1), HuCoV-NL63 (ABI20791.1), HuCoV-229E (N protein_073556.1), HuCoV-HKU1 (AYN64565.1), HuCoV-OC43 (QBP84763.1) were obtained from GenBank for sequence alignments. For generation of *E. coli* expression plasmids, amino acid sequences from each virus (SARS-CoV-2 (132-419), SARS-CoV (133-422), HuCoV-NL63 (100-377), and HuCoV-229E (102-389), were used to generate bacterial codon optimized DNA gBlocks using the Integrated DNA Technologies web server tool. A list of gBlocks and PCR primers can be found in Table S1. Each gBlock was cloned into the pET28a T7 expression vector using the New England Biolabs NEBuilder Assembly 2X Master Mix according to the manufacturer’s instructions. The resulting clones (SARS-CoV-2 = pBG690 | SARS-CoV = pBG700 | HuCoV-NL63 = pBG702 | HuCoV-229E – pBG705) were sequence verified by Sanger sequencing (Genewiz).

Nucleocapsid proteins were expressed in BL21 DE3 pLys *E. coli* and purified by nickel affinity and gel filtration chromatography essentially as previously described (26) with the exception of using 25 mM HEPES (pH 7.5) as the buffer and changing the final NaCl concentration to 500mM in all of the buffers to reduce oligomerization of N proteins. N proteins were concentrated to ∼2mg/ml in gel filtration buffer (25 mM Hepes pH 7.5, 500 mM NaCl, 1 mM DTT, 0.1 mM AEBSF, and 10% glycerol) using 10K Centricon concentrators, flash frozen in liquid nitrogen, and stored at −80°C until use.

### N protein generation and initial screening of monoclonal antibodies

All immunizations were intraperitoneal in a final volume of 200μl containing 25 ug of recombinant N protein antigen. Two 6-week old female BALB/c mice (Jackson Laboratories) were primed with antigen emulsified in an equal volume of complete Freund’s adjuvant (Millipore-Sigma). The mice were boosted 2 and 4 weeks later with antigen emulsified in incomplete Freund’s adjuvant. Mice were bled from the tails at 2 and 6 weeks to verify seroconversion by ELISA. A final boost of antigen in PBS at 8 weeks followed by euthanasia and spleen retrieval 4 days later. On the day of splenocyte collection, immunized mice were euthanized by CO_2_ asphyxiation followed by exsanguination. Mouse carcasses were sterilized with 70% ethanol before the spleen was removed and eviscerated. After passing through a 16-gauge needle, the splenocytes were washed and fused with Sp2/0 Ag14 myeloma cells using the ClonaCell™-HY Hybridoma Kit. Following fusion, hybridoma cells were rested for 24 hours before being resuspended and plated in ten 10cm plates with ClonaCell™-HY Semi-Solid Medium D and allowed to propagate for 10 days at 5% CO_2_ at 37°C. Eleven days after fusion, individual colonies were selected and transferred to individual wells in a 96-well plate using a 10μl micropipetter. 920 colonies were harvested and left to grow for five days at 5% CO_2_ and 37°C.

### Primary Hybridoma Screen

Hybridomas were screened using enzyme-linked immunosorbent assay (ELISA) for activity against SARS-CoV-2 N protein. 96-well plates were coated with 2ug/ml N protein diluted in 1X PBS and incubated overnight at 4°C. Plates were blocked with SuperBlock™ T20 (TBS) for 1 hour shaking at room temperature. After three washes with 0.1% Tween in 1X PBS (Hyclone), supernatant from the 920 hybridoma colonies were incubated on the plates for 1 hour shaking at room temperature. After three more washes the plates were incubated with HRP conjugated goat anti-mouse polyclonal antibodies (Abcam ab97023) diluted at 1:10,000 in 1X PBS for 1 hour at room temperature, shaking. The plates were then washed three more times before being developed with 3,3’,5,5’-Tetramethylbenzidine (TMB) before the reaction was stopped with an equal volume of 2M H_2_SO_4_. Absorbance at 450nm was determined for each well using a PerkinElmer Victor X5 multilabel plate reader. Absorbances were corrected against the PBS negative control and organized by absorbance on Microsoft Excel. Ninety-two colonies with the highest absorbance at 450nm were selected for further testing.

### Bacteria Cross-Reactivity Screen

One 96-well plate was coated with 200ng/well of recombinant N protein and another plate was coated with 200ng/well of *E. coli* lysate (BL21 DE3 pLys). The aforementioned ELISA protocol was performed using the selected 92 parent hybridoma culture supernatants as the primary antibody. Following A_450_ determination any colonies with reactivity towards the bacterial lysate protein excluded from further screening.

### His-Tag Cross-Reactivity Screen

One 96-well plate was coated with 200ng/well of recombinant N protein and another plate was coated with 200ng/well of recombinant 6xHis-tagged SARS-CoV-2 spike protein receptor binding domain (RBD) produced using the FreeStyle 293 Expression System (Thermo Fisher) from BEI Resources (Cat# NR-52366) as described (27). The ELISA protocol described previously was performed using the selected 92 parent hybridoma culture supernatants as the primary antibody. Following A450 determination and correction in relation to the Medium E negative control, any colonies with reactivity towards the His tags were removed from consideration.

### SARS-CoV-2 virus

SARS-CoV-2 coronavirus (Isolate USA-WA1/2020) was obtained from BEI Resources. Stocks of virus were grown in Vero-E6 cells in DMEM (Gibco) supplemented with 10% fetal bovine serum (Atlas Biologicals) and 25mM HEPES (pH 7.5) in BSL-3+ containment at the Colorado State University Regional Containment Biocontainment Laboratory. Virus containing media was stored at −80°C in single use aliquots. Viral titers were performed using plaque assay as described (28).

### Immunofluorescence assay

Vero cells were plated at a concentration of 3.5×10^4^ cells/well in a 96-well plate. The cells were inoculated with SARS-CoV-2 in BSL-3 at a MOI of 0.1 and allowed to absorb for 1 hour at room temperature. Unabsorbed virus was washed with 1X PBS, cells were overlaid with 1X DMEM supplemented with 2mM glutamine, non-essential amino acids and 2% fetal bovine serum, and the cells were incubated for 24 hours at 5% CO_2_ and 37°C. Infected cells were briefly washed with 1X PBS then fixed with either 4% paraformaldehyde (PFA) or methanol for 10 minutes before being washed three more times. PFA fixed cells were permeabilized by incubating with 0.5% Tween in 1X PBS for 20 minutes.

Plates of infected fixed cells were blocked with 4% nonfat dry milk powder dissolved in 1X PBS + 0.1% Tween 20 for 1 hour at room temperature, shaking. Supernatant from parents 17, 21, 22, 57, and 67 along with rabbit anti-N protein polyclonal control antibody were then incubated for 1 hour shaking at room temperature and then washed three times with 1X PBS + 0.1% Tween 20. FITC-conjugated goat anti-mouse secondary antibodies or Alexa 488-conjugated goat anti-rabbit secondary antibodies at a dilution of 1:2000 in 1X PBS were added to the appropriate wells, with all wells receiving Hoescht staining at 1:4000. After 1 hour of incubating the secondary antibodies and stains in the dark at room temperature, the plates were washed three times with 1X PBS + 0.1% Tween 20 with the last wash being 50μl of 1X PBS. Cell labeling and fluorescence were observed with a Celigo high-content imaging cytometer (Nexcelcom) and a Nikon Diaphot 200 fluorescence microscope.

### Western Blot

The ability for the parent colonies’ antibodies to detect linear epitopes was assessed via western blotting. Recombinant N protein and recombinant spike RBD proteins were resolved on a 12% gel in triplicate at 120V for 1 hour before being transferred to PVDF membranes for one hour at 100V. Blots were blocked with 4% nonfat dry milk powder blocking solution in 1X PBS. Supernatants were diluted 1:5 in blocking solution (PBS+2% non-fat dry milk) before being applied to blocked blots overnight, shaking at 4LC. Blots were washed three time with blocking solution for five minutes each before being incubated for 1 hour at room temperature with goat anti-mouse HRP-conjugated secondary antibodies diluted in blocking solution. After three more washes with 1X PBS, the blots were developed with 1-Step™ Ultra TMB Blotting Solution (Pierce) before being quenched in deionized water and imaged.

Reactivity towards endogenously produced N protein from an active SARS-CoV-2 infection was assessed with infected cell lysates on western blots. Vero cells were infected at 0.1 MOI with SARS-CoV-2 in BSL-3 and allowed to incubate for 48 hours before cells were trypsinized, spun down, and resuspended in RIPA buffer. These samples were then diluted in 2X Laemmli Buffer and boiled for 15 minutes before being resolved on an SDS-PAGE gel alongside uninfected Vero cell lysate and processed for western blot analysis as described above.

### Isotyping

The isotype of the antibodies produced by parent hybridomas was assessed using the Pierce™ Rapid Antibody Isotyping Kit plus Kappa and Lambda (catalog no. 26179) for mouse antibodies according to the provided procedure. Briefly, antibodies were diluted 1:100 in the provided sample diluent before being applied to the lateral flow assay and the corresponding bands observed after 10 minutes.

### Sequencing of anti-N protein monoclonal antibody genes

RNA from each hybridomas 17, 21, 22, 57, and 67 was isolated and stored at −80°C following Trizol (Invitrogen) and phenol-chloroform extraction. Isolation of kappa, lambda, and heavy chain RNA sequences for hybridoma colonies was accomplished using the monoclonal antibody sequencing protocol established in Meyer et al. 2019 (29). Primers specific for the 3’ constant region of either the kappa, lambda, or heavy chain RNA sequences outlined in the protocol were used in conjunction with SMARTScribe Reverse Transcriptase. Amplification of the RT product is accomplished with PCR using primers specific to the universal sequence and a region of the kappa, lambda, or heavy chain sequences that were offset to the primers used during reverse transcription. Successfully produced PCR products were then cloned into the NEB pMiniT 2.0 *Escherichia coli* vector using the NEB PCR Cloning Kit. Five kappa and heavy chain clones were sequenced for each hybridoma by Genewiz. Clone sequences were analyzed by IgBLAST (30) for antibody framework regions (FMR) and complementary-determining regions (CDR).

### Epitope Mapping

Primers described in Table S1 in were used to create 50 amino acid deletions from the pBG690 plasmid used to produce SARS-CoV-2 N protein amino acids 133-419. The primers produced recombinant N protein variant sequences corresponding to proteins Δ133-179, Δ180-229, Δ230-279, Δ325-379, and Δ381-419. These transcripts were circularized by NEBuilder and then cloned into BL21 *E. coli* cells. Protein expression was incubated overnight in LB broth with 10% glucose. Recombinant N protein production was then induced with IPTG alongside uninduced transformed variants for five hours. Cells were then spun down and resuspended in Laemmli Sample Buffer and boiled for five minutes and then run on a 10% SDS-PAGE gel. Protein was then transferred to a PVDF membrane and then labeled according to the aforementioned western blot protocol before being developed with TMB.

## Results

This project was focused on developing novel antibodies that are able to specifically detect SARS-CoV-2 nucleocapsid protein for use in research and diagnostic testing efforts. Therefore, recombinant N protein was produced as antigen for hybridoma production. The N-terminal domain of human coronavirus N proteins have several conserved regions that may contribute to monoclonal cross-reactivity (**Fig. 1A**). Previous work developing monoclonal antibodies against MERS coronavirus N protein found that removal of the N-terminal domain improved antibody specificity and increased recombinant protein solubility (31). Therefore, we developed a bacterial expression plasmid that produces AA133-416 of the SARS-CoV-2 (isolate USA-WA1/2020) N protein with a N-terminal T7 leader sequence to improve translation efficiency (pBG690). Expression of recombinant N protein in BL21 DE3 pLys E. coli was relatively robust, but we found that the recombinant protein formed large molecular weight oligomers at NaCl concentrations below 300 mM (data not show). Purification of nickel-affinity purified N protein on a Superdex 200 gel filtration column in 500 mM NaCl resolved these aggregation issues and produced protein migrated as the predicted molecular weight of a truncated N dimer (**Fig. 1B**). SDS-PAGE analysis of purified SARS-CoV-2 N protein showed that the protein was >98% pure and migrated at the expected molecular weight (**Fig. 1C**). This purified antigen was found to be antigenic and produced rabbit polyclonal antibodies that reacted with N protein in deer mouse infections in parallel work (32). The recombinant N protein was used to immunize BALB/c mice according to the protocol described above to generate immunized mice, from which spleens were collected and hybridoma clones were produced.

**Figure 1.**
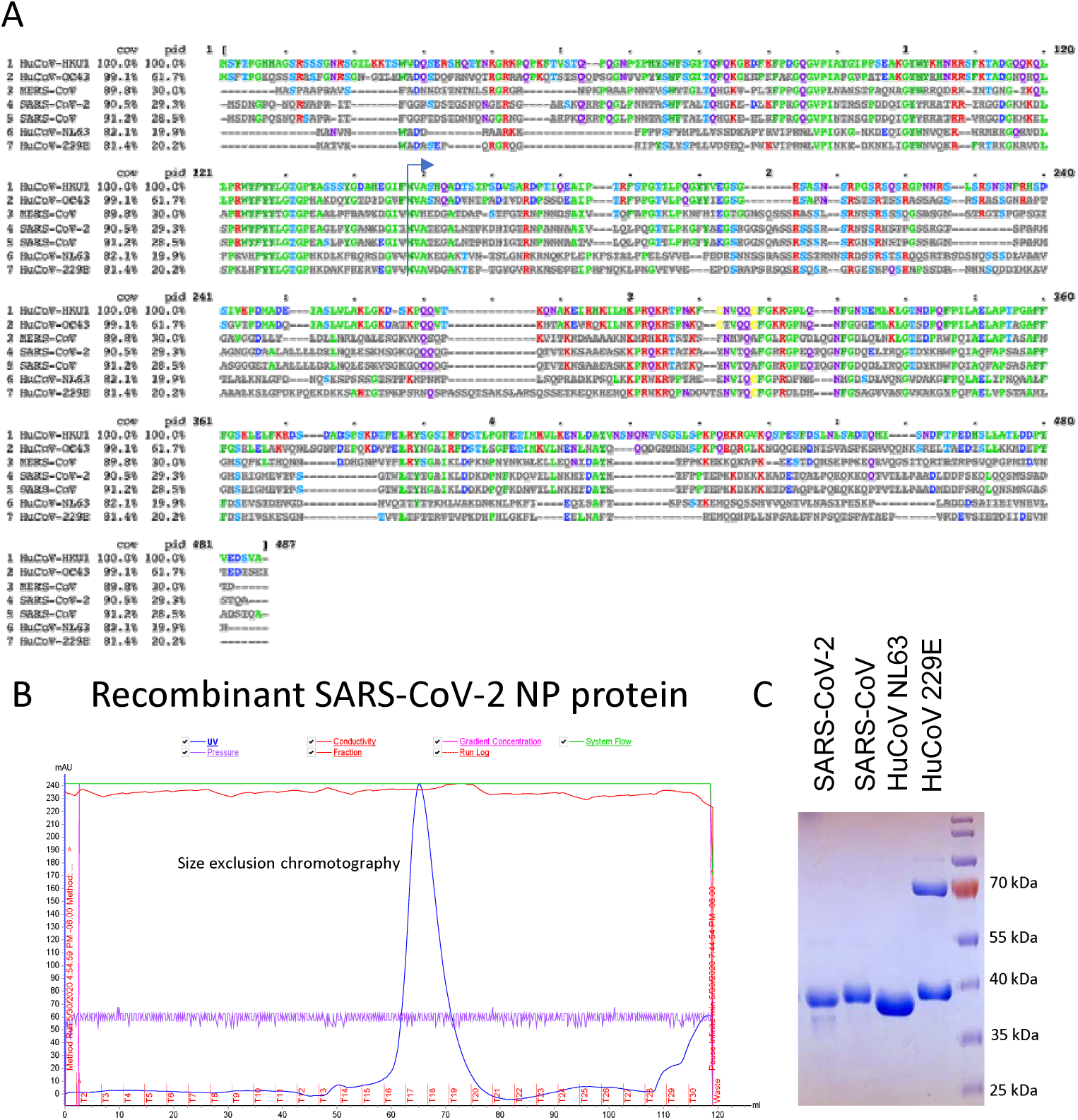
Production and characterization of truncated SARS-CoV-2 nucleocapsid protein. **A**) Sequence alignment of human coronavirus N proteins. **B)** Size exclusion chromatograph of nickel-column purified SARS-CoV-2 NP (133-419).**C)** SDS-PAGE gel of purified Coronavirus nucleocapsid proteins.

For primary screening of hybridoma colonies, supernatant was collected from the 920 colonies and tested for the presence of N protein specific antibodies by ELISA (**Fig. 2A**). After determining reactivity of produced monoclonal activities against the antigen via light absorbance at 450nm and correcting based on the negative control, the highest scoring 92 hybridoma colonies were selected for further testing and expansion as “parent” colonies. Due to the bacterial system used for N protein immunogen production, we then screened these 92 parents for reactivity against bacterial proteins. Plates were coated with bacterial lysate protein (BLP) and tested by ELISA in parallel to N protein-coated plates to determine which parents had residual bacterial reactivity. Of these, parents 5,7, 23, 43,53, 54, 65, 77, and 79 demonstrated specificity for bacterial proteins and were thus eliminated from further testing. The remaining parents were then tested for reactivity against the 6xHis tag modification attached to the N protein immunogen and antigen used during the primary screen. Parent 55 showed reactivity to the 6xHis tag and was thus eliminated from further testing. After assessing the reactivity towards N protein in the primary screen, BLP screen, and His tag screen in contrast to reactivity towards BLP and His, 18 parents were selected to be top parents for further testing and given the prefix mBG (**Fig. 2B**). Later in the project we produced recombinant protein for SARS-CoV, HuCoV-NL63, and HuCoV-229E N proteins (**Fig. 1C**), but these proteins were not available until late in the hybridoma screening process and could not be used to counter-screen hybridomas during the initial screening process.

**Figure 2.**
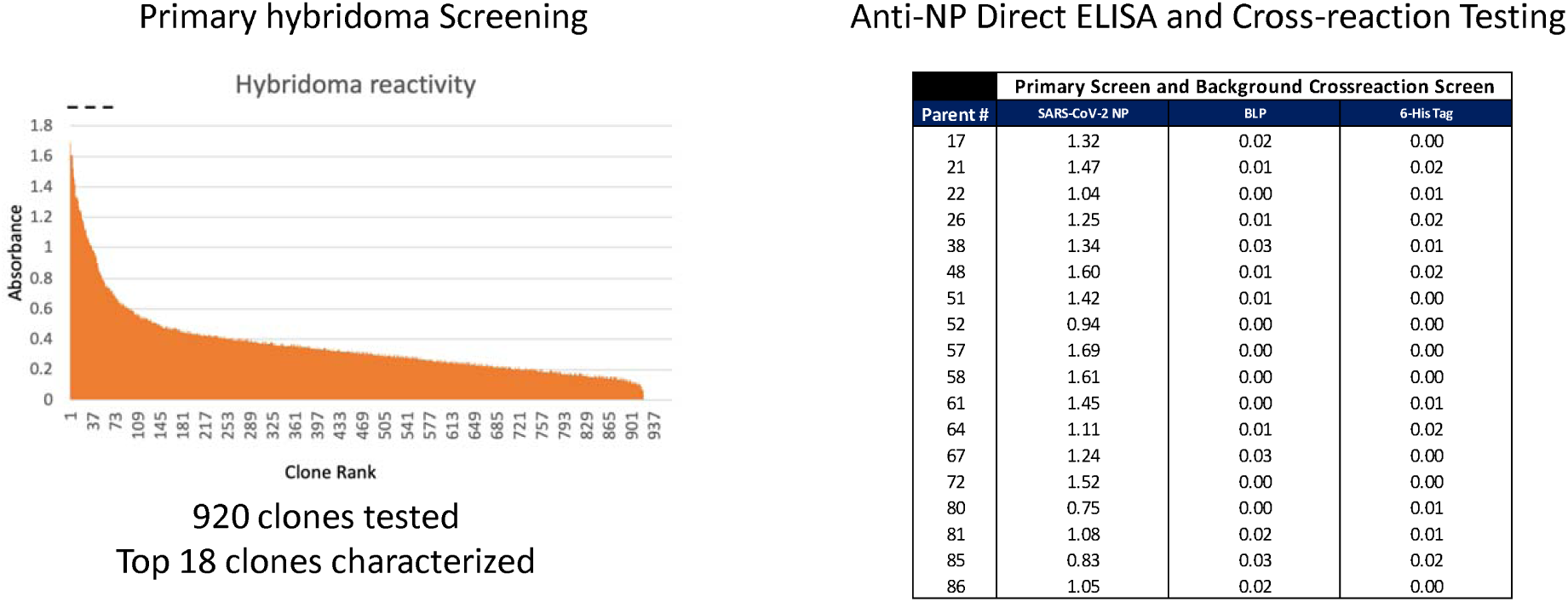
Screening of Anti-Nucleocapsid Clones. **A)** Direct ELISA analysis of 920 clones picked from hybridoma fusion. **B)** Verification ELISA of top 18 anti-nucleocapsid monoclonal antibody clones and counter screening against bacterial lysate (BLP) and 6-His-tagged SARS-CoV-2 Spike RBD domain. Averages are presented following background subtraction.

Recombinant SARS-CoV N protein, HuCoV-NL63 N protein, and HuCoV-229E N protein were expressed and purified for determining the top 18 parents’ specificity compared to SARS-CoV-2 N protein. After ELISA testing and analysis, the parents’ reactivity towards these variants of N protein were compared (**Table 1**). With the exception of mBG 61, 64, and 86 there was strong cross reactivity with SARS-CoV N protein. In a diagnostic or research environment, such cross reactivity is of limited concern due to the low prevalence of SARS-CoV circulating in communities. Cross reactivity with NL63 N protein was similar in that mBG 61, 64, and 86 lacked reactivity to this N protein variant while other parents maintained some reactivity, albeit much less than they demonstrated for SARS-CoV. mBG 21, 22, 57, and 67 all had reduced reactivity towards NL63 N protein. All parents lacked reactivity towards 229E N protein, whereas the anti-His control showed equal reactivity towards all N proteins tested.

**Table 1.**
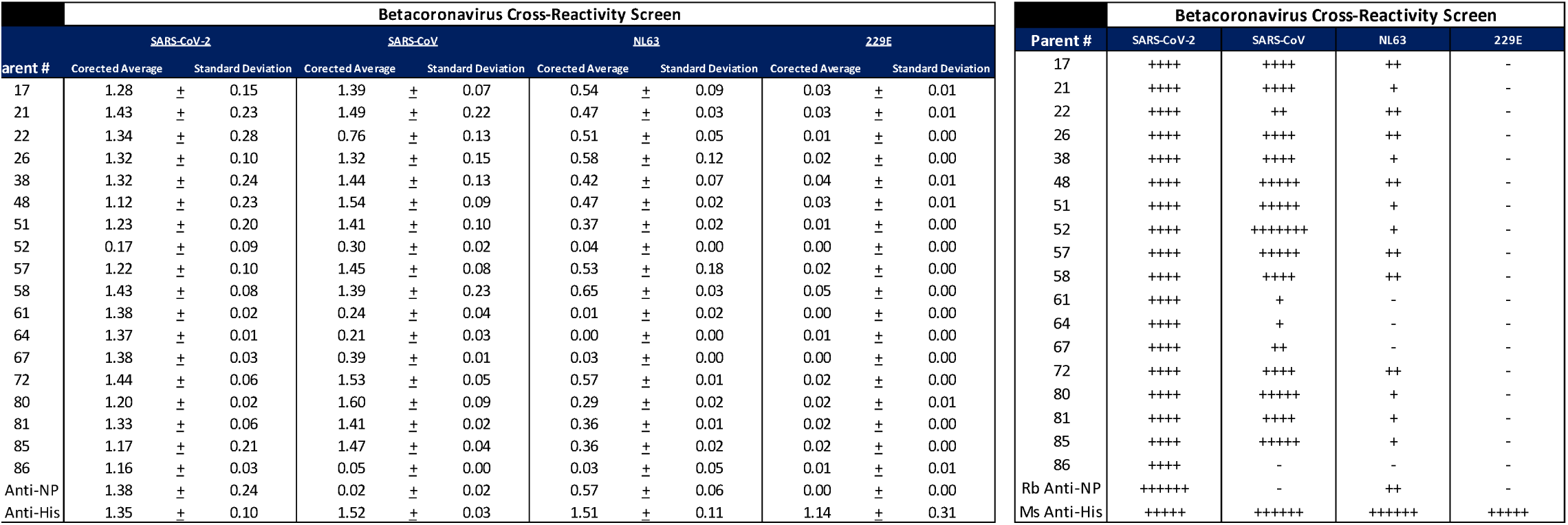
Cross-Reactivity Screening of Anti-Nucleocapsid Monoclonal Antibody Clones. **Left:** Direct ELISA analysis of top 18 clones against SARS-CoV2, SARS-CoV, NL63, and 229E human coronavirus nucleocapsid proteins. Average signals are corrected against background signal **Right:** Relative binding of monoclonal antibodies to nucleocapsid proteins compared to SARS-CoV-2.

Western blots were performed to determine reactivity of the top 18 parents towards linear or semi-linear epitopes. All parents demonstrated specific reactivity for recombinant SARS-CoV-2 N protein (**Supplemental Fig. 1**) with any residual bands in the neighboring wells being the result of bleed-over of sample during loading. The multiple bands on each blot corresponds to oligomers of N protein and degraded protein. mBG 67, 80, and 86 demonstrated two band labeling each, suggesting a specificity for linear epitopes on the N protein while the other parents labeling suggests conformational epitopes. Western blots comparing reactivity towards SRS-CoV-2 infected Vero cell lysates and uninfected Vero cell lysates (**Fig. 3**) demonstrated a high specificity towards antigen in infected cells and the endogenously produced N protein for the top 18 clones.

**Figure 3.**
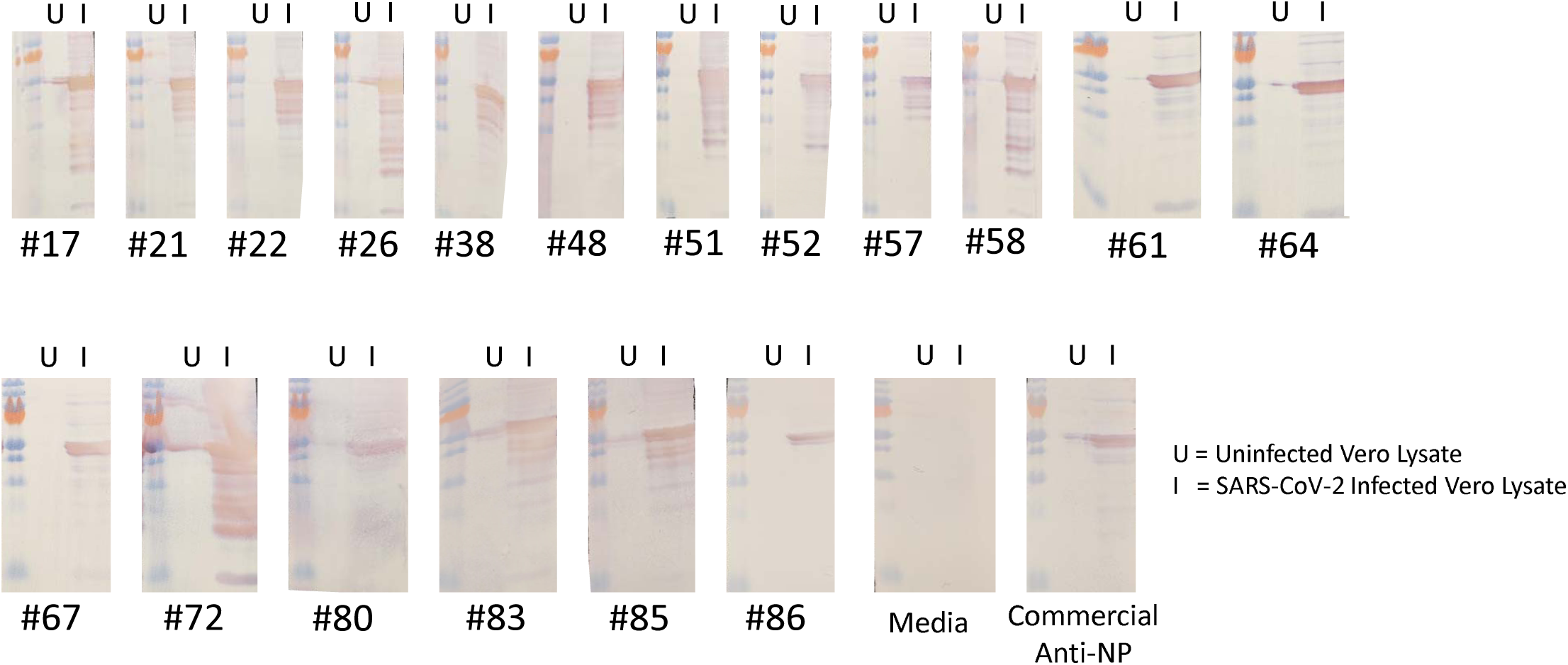
Western Blot Analysis of Anti-Nucleocapsid Monoclonal Antibody Clones Against Uninfected (U) or SARS-CoV-2 infected (I) Vero cells.

To test the capability of these antibodies to be used for cellular localization of N in SARS-CoV-2 infected cells, the ability for the top 18 parents to label endogenous N protein following PFA or methanol fixation was determined by immunofluorescence assay (IFA). Signals intensity and subcellular localization were comparable to a previously described rabbit anti-NP polyclonal antibody (32) and **Fig. S2**. Results indicated that the parents’ antibodies that demonstrated reactivity to fixed infected cells were non-reactive towards uninfected fixed cells. N protein was found predominantly in the cytoplasm in SARS-CoV-2 infected cells (**Fig 4B**). IFA positive clones varied in effectiveness between methanol and paraformaldehyde fixation methods (**Fig. 4A**).

**Figure 4.**
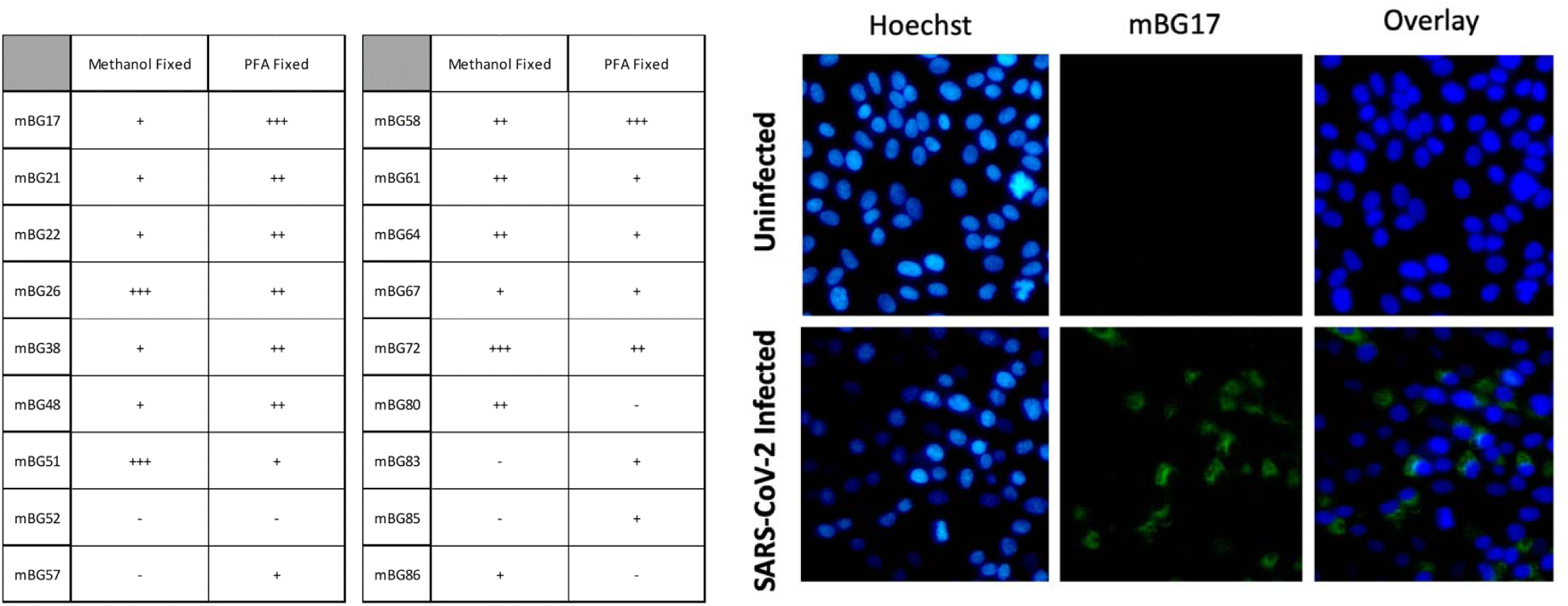
Immunofluorescence Analysis of Anti-Nucleocapsid Monoclonal Antibody Clones. **A)** Relative reactivities of clones in SARS-CoV-2 infected Vero cells fixed with methanol or paraformaldehyde. **B)** representative images of uninfected and SARS-CoV-2 infected Vero cells (paraformaldehyde fixed) processed for immunofluorescence analysis with mBG17.

Based on the activity of clones in ELISA, western blot, and IFA analysis, we chose five clones (mBG17, mBG21, mBG22, mBG57, and mBG67) for sequence analysis. Sequence determination via the method outline in Meyer et al., with five separate clones for each V_H_ or V_L_ being sequenced (29). Consensus sequences for each clone were assembled, translated, and subjected to IgBLAST analysis to determine framework and complementary determining regions (**Table 2**). mBG21 and 22 presented identical sequences, suggesting a common B-cell origin. Isotype tests revealed each of the five parents, with the identical isotypes of mBG 21 and 22 supporting the prediction of a common origin. Clones mBG21/22, and mBG57 shared identical heavy chains but different kappa chains. Clones mBG17 and mBG67 possessed unique heavy and light chains.

**Table 2.**
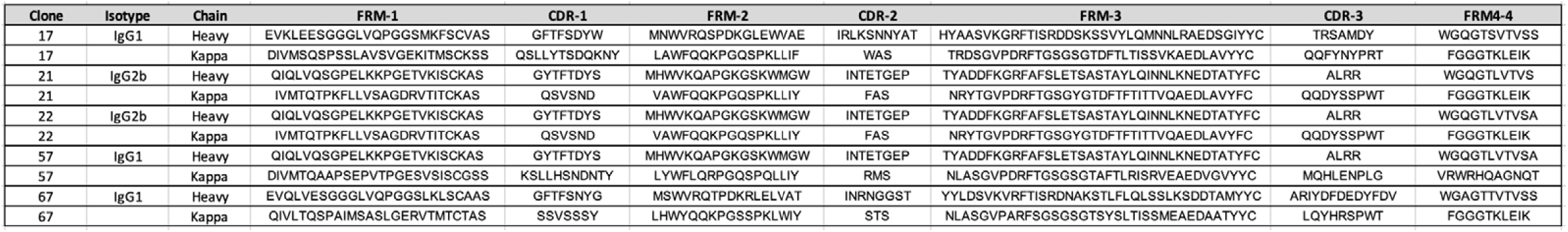
Immunoglobulin heavy and light chain amino acid sequences of top 5 hybridoma clones (as defined by IgBlast).

Finally, we sought to determine preliminary epitope ranges for clones mBG17, mBG21/22, mBG57, and mBG67 using SARS-CoV-2 N protein deletion mutants in western blot analysis. Recombinant N protein expression plasmids, each with a different 50 amino acid deletion, were constructed and used to narrow down the range of epitope locations for mBG17, mBG22, mBG57, and mBG67 (**Fig. 5**). Clones mBG22, mBG57, and mBG67 showed strong reactivity in western blot analysis against all clones except the Ll133-179 deletion, suggesting that mBG21/22, mBG57, and mBG67 epitope resides within AA133-179 range. mBG17 showed strong reactivity towards all deletions except Δ381-419, indicating that clone mBG17 is likely within AA381-419 at the C-terminal end of the N protein. Additional peptide mapping is needed to determine the exact amino acid sequences for each epitope.

**Figure 5.**
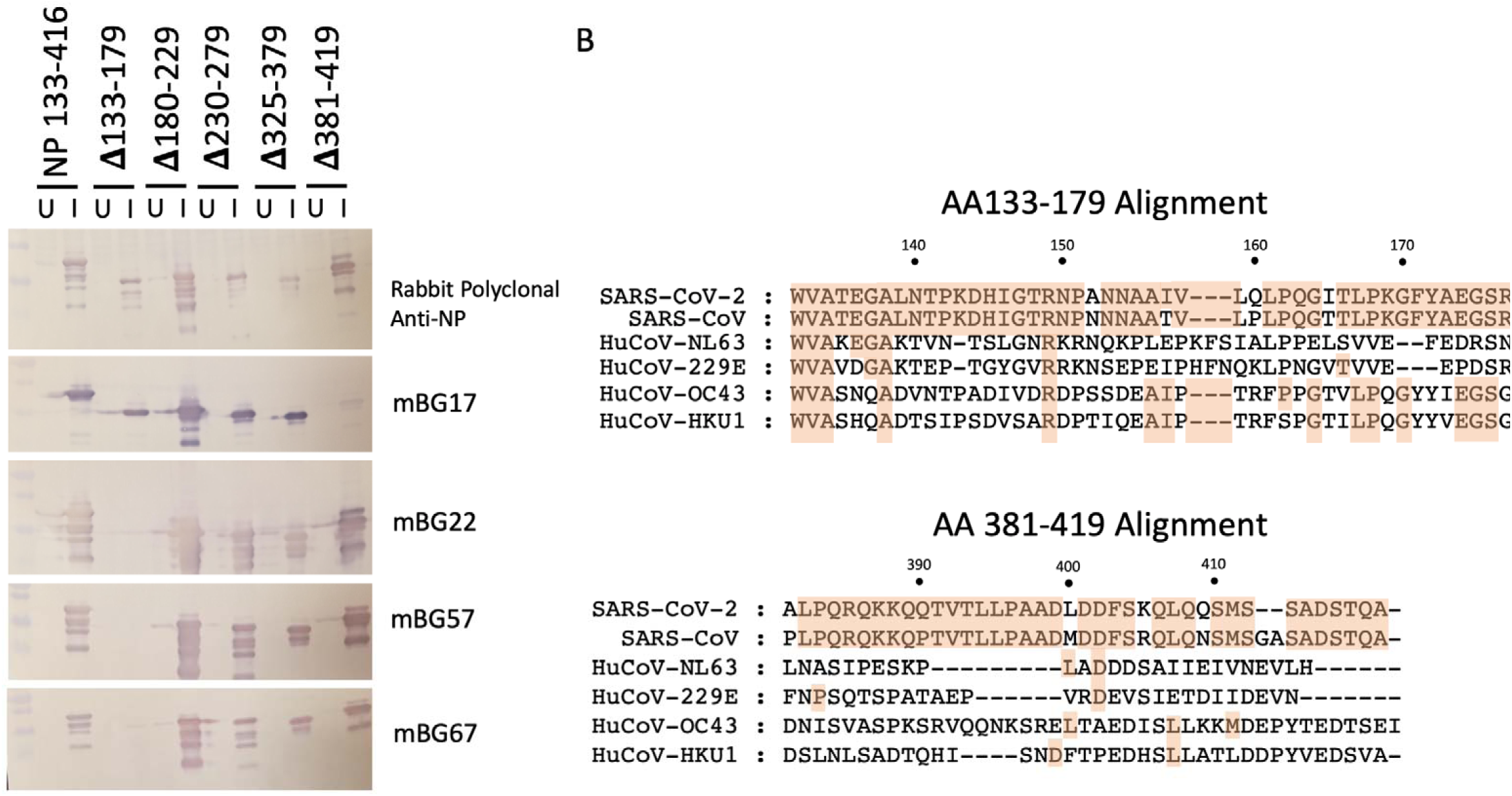
Epitope mapping using N protein deletions. **A)** Western blot analysis of selected antibody reactivity against SARS-CoV-2 nucleocapsid protein deletions. **B)** Sequence alignments of NP AA133-179 and AA381-419 regions with heterologous human coronavirus nucleocapsid proteins.

## Conclusions

With SARS-CoV-2 spreading globally amidst a dearth of effective diagnostics, treatments, and reagents to tackle pandemic, it is more important than ever for tools for detection and research are developed and validated. The hybridoma antibodies characterized in this paper have been selected and validated for high specificity towards SARS-CoV-2 nucleocapsid protein across several diagnostic and research assays, such as direct ELISA, IFA, and western blot. Future studies can utilize these antibodies for studies determining N protein structure, intracellular interactions, diagnostic development, and potential therapeutics. Preliminary epitope mapping experiments revealed ranges of amino acids that contain partial or full epitopes utilized by antibodies from five of the top mBG antibodies. The sequences for the FMR and CDR regions of these antibodies shown in **Table 2** be used for the production of recombinant single-chain variable fragment (scFv) antibodies that may be used for observing real-time N protein production (33), heavy and light genes cloned into antibody expression vectors for non-hybridoma eukaryotic expression systems, or used in neutralizing treatments (34). Overall, we hope that these data will be useful to the wider research community for fighting the ongoing COVID-19 pandemic.

## Acknowledgements

We would like to thank the members of the Geiss, Perera, and Schountz labs for helpful discussions. We would also like to thank Jeff Wilusz for assistance with monoclonal antibody production. This work was funded in part by an NIH grant (R01 AI132668) to BJG, NIH grant (R01 AI140442) to TS, The Boettcher Foundation COVID Biomedical Research Innovation Fund award to RP and by support from the Colorado State University Department of Microbiology, Immunology and Pathology and Colorado State University Office for the Vice President of Research.

## Figure Legends

**Supplemental Figure 1. Western blot analysis of anti-SARS-CoV-2 N protein monoclonal antibodies against recombinant antigen**

**Supplemental Figure 2. IFA images of SARS-CoV-2 infected and uninfected cells stained with control rabbit anti-NP polyclonal antibody. Images obtained with a Celigo high-content imaging cytometer (Nexcelcom)**

**Supplemental Table 1. DNA gBlocks and oligonucleotide primers**

