## Supplemental Data for "Development of SARS-CoV-2 Nucleocapsid Specific Monoclonal Antibodies"

### Slide 1
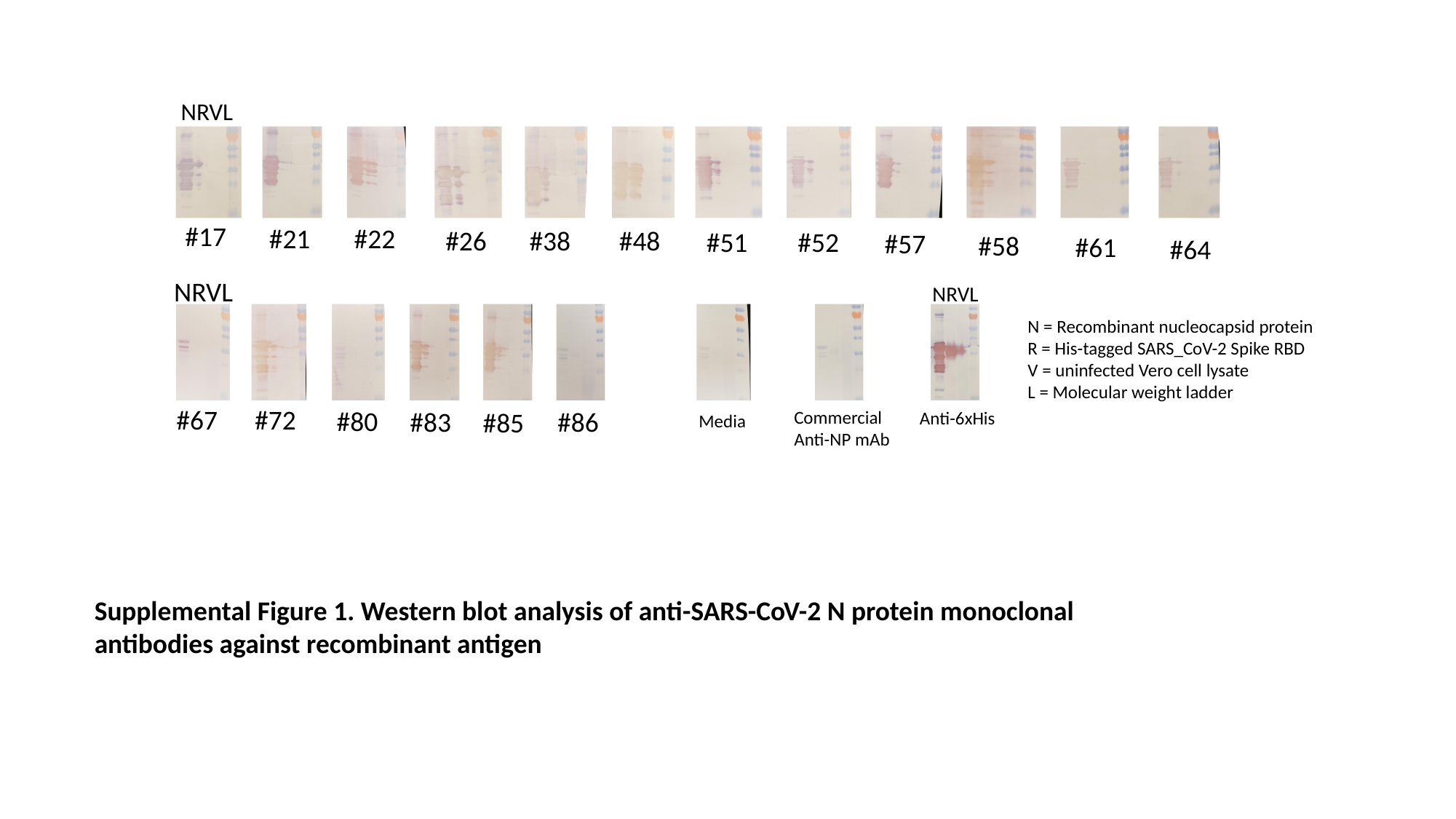

NRVL
#17
#21
#22
#26
#38
#48
#51
#52
#57
#58
#61
#64
NRVL
NRVL
N = Recombinant nucleocapsid protein
R = His-tagged SARS_CoV-2 Spike RBD
V = uninfected Vero cell lysate
L = Molecular weight ladder
#67
#72
#80
#86
#83
#85
Commercial
Anti-NP mAb
Anti-6xHis
Media
Supplemental Figure 1. Western blot analysis of anti-SARS-CoV-2 N protein monoclonal antibodies against recombinant antigen

### Slide 2
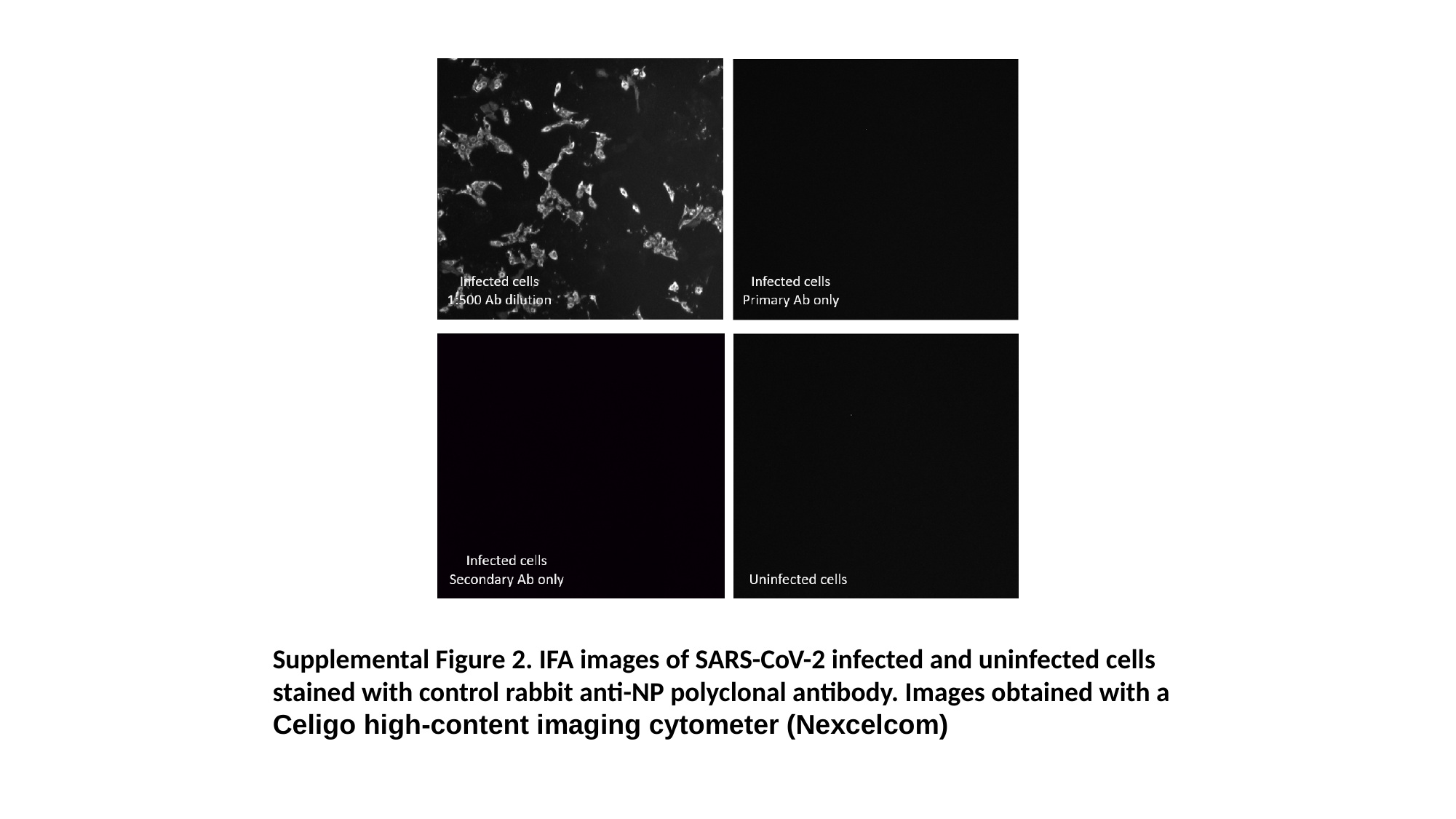

Supplemental Figure 2. IFA images of SARS-CoV-2 infected and uninfected cells stained with control rabbit anti-NP polyclonal antibody. Images obtained with a Celigo high-content imaging cytometer (Nexcelcom)

### Slide 3
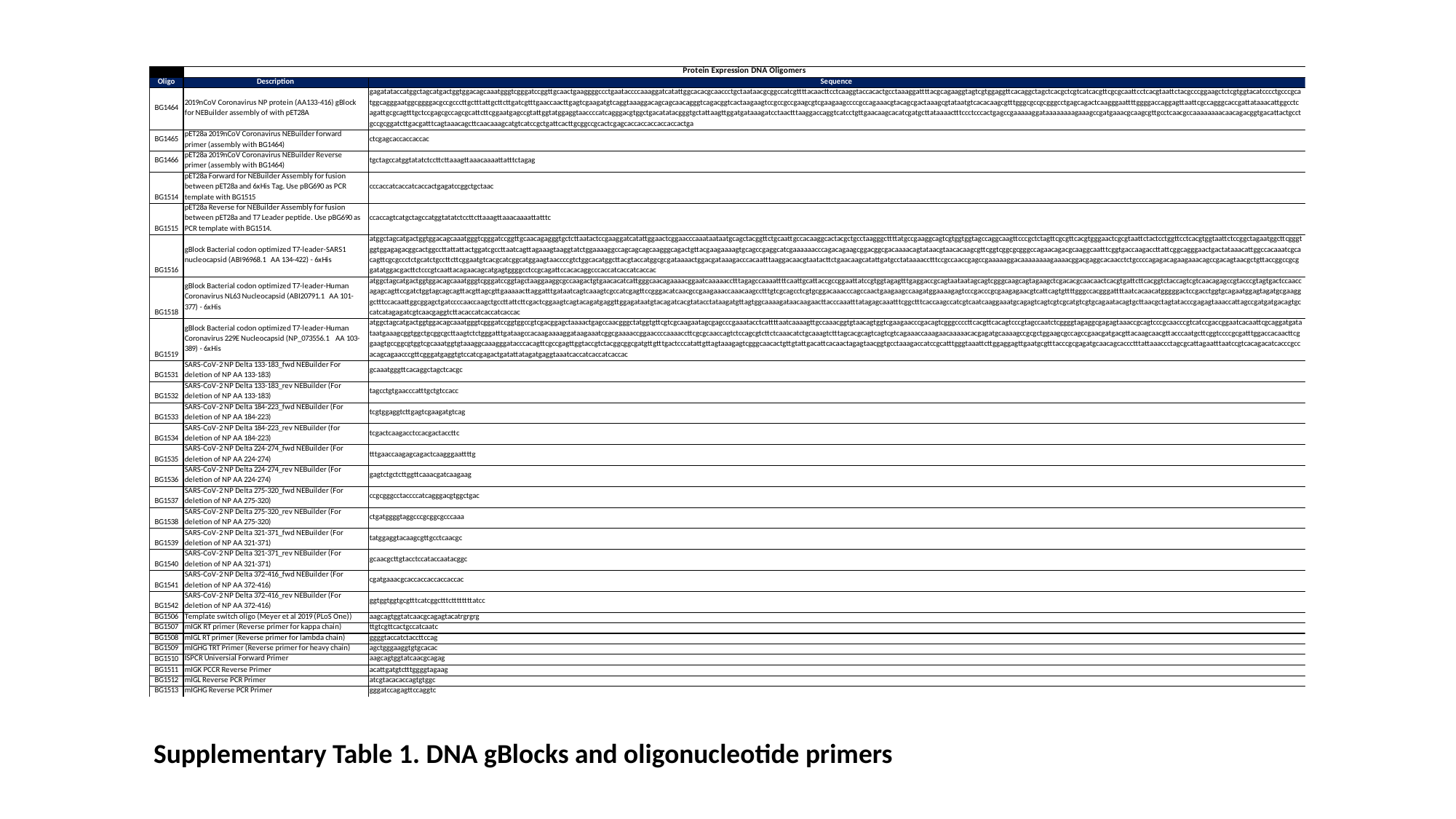

Supplementary Table 1. DNA gBlocks and oligonucleotide primers
